# Insights into the mechanism of bovine spermiogenesis based on comparative transcriptomic studies

**DOI:** 10.1101/2020.09.25.313908

**Authors:** Xin Li, Chenying Duan, Ruyi Li, Dong Wang

## Abstract

To reduce the reproductive loss caused by semen quality and provide theoretical guidance for the eradication of human male infertility, differential analysis of the bovine transcriptome among round spermatids, elongated spermatids, and epididymal sperm was carried out with the reference of the mouse transcriptome, and the homology trends of gene expression to the mouse were also analysed. First, to explore the physiological mechanism of spermiogenesis that profoundly affects semen quality, homological trends of differential genes were compared during spermiogenesis in dairy cattle and mice. Next, the Gene ontology (GO), Kyoto Encyclopedia of Genes and Genomes (KEGG) pathway enrichment, protein-protein interaction network (PPI network), and bioinformatics analysis uncovered the regulation network of acrosome formation during the transition from round to elongated spermatids. In addition, processes that regulate gene expression during spermiogenesis from elongated spermatid to epididymal sperm, such as ubiquitination, acetylation, deacetylation, glycosylation, and the functional gene ART3 may play an important role during spermiogenesis. Therefore, its localisation in the seminiferous tubule was investigated by immunofluorescent analysis, and its structure and function were also predicted. This study provides important data for revealing the mystery of life during spermiogenesis resulting from acrosome formation, histone replacement, and the fine regulation of gene expression.

## Introduction

With the extensive application of artificial insemination, breeders have progressively focused on semen requirements for the quality and quantity of excellent bulls. Approximately 10 to 60 billion sperms can be obtained from bulls when the semen is collected at a frequency of three times a week (1). Considering the presence of 20 million sperms per straw and a calving rate of 53% (2), each bull can theoretically breed about 13,000 to 78,000 offspring yearly. However, due to semen quality issuses caused by insufficient spermiogenesis, the pregnancy rate of each straw of frozen semen varied by about 20% (2), resulting in a tremendous waste. At the same time, about 15% of the couples of childbearing age worldwide are affected by infertility, of which 50% are due to male factors (3), and even sperm with normal morphology can cause infertility (4). Therefore, it is necessary to explore the mechanism of spermiogenesis, identify important regulatory pathways and important functional genes to improve the semen quality and yield of herd sire, and overcome the problem of male infertility (2,5).

Since histones are gradually replaced by protamine in the process of spermiogenesis, chromosomes are highly condensed, resulting in the overall shrinkage of the nucleus. At the same time, the spermatid gradually polarises, and the Golgi bodies progressively aggregate and transfer to one end of the spermatid, and finally become specialised into the acrosome, forming the sperm head together with the nucleus. The centrioles move in the direction opposite to the head and extend outward to become the skeleton of the sperm tail, while mitochondria gradually specialise to form a ring-shaped mitochondrial sheath, which is attached to the outside of the axon filament in the midpiece of the sperm tail. At the same time, the cytoplasmic residue gradually fell off. Following these changes, the round spermatid eventually develops into the tadpole-like sperm with head and tail, which are mature and stored for a short time in the epididymis (Figure 1).

**Figure 1.**
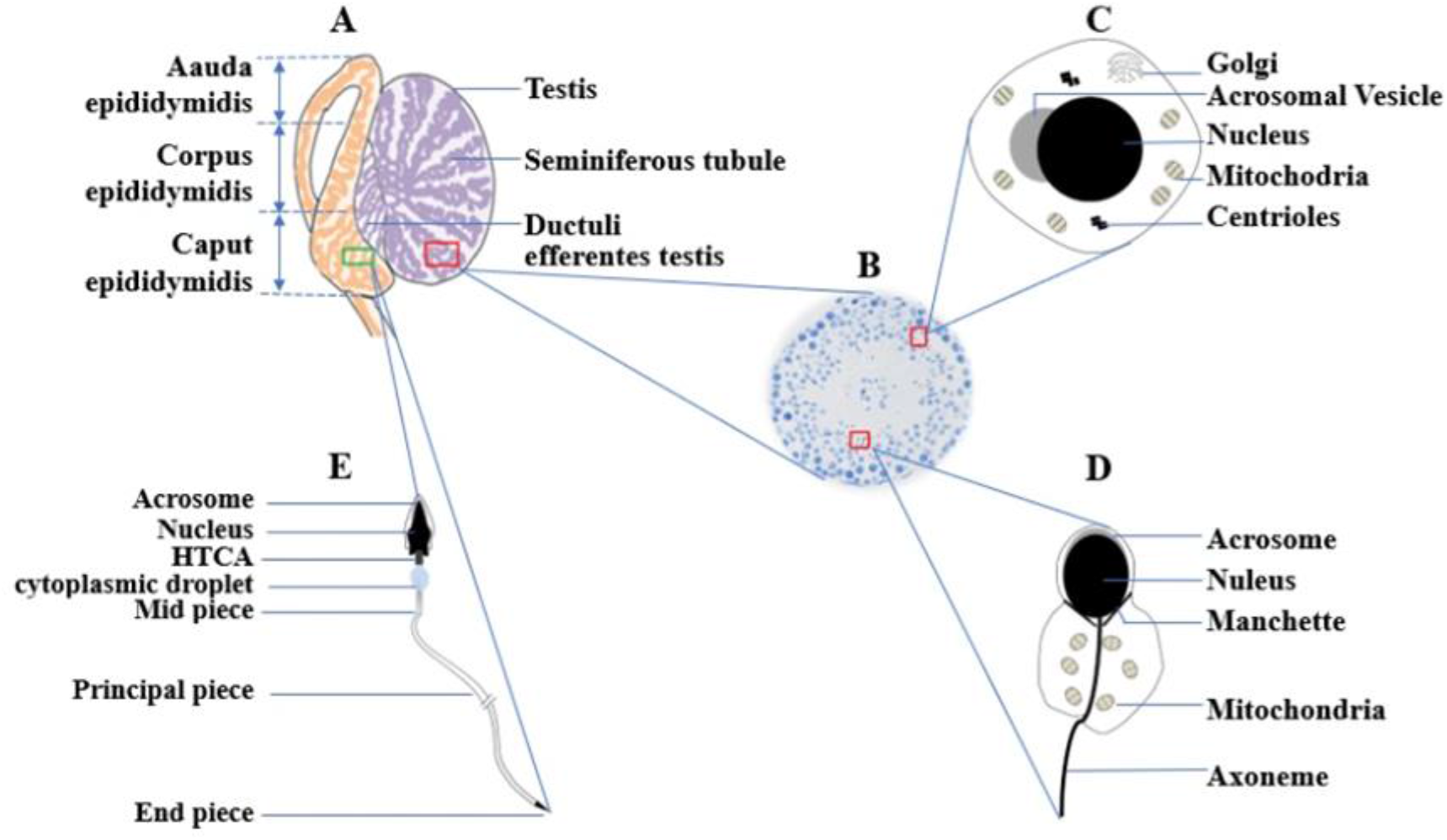
A representative schematic of spermiogenesis. (A) Testis and epididymis. (B) The cross section of seminiferous tubule. (C) Round spermatid. (D) Elongated spermatid. (E) epididymal sperm. *HTCA* stands for the head tail coupling apparatus.

However, abnormal expression of some important functional genes during spermiogenesis affects the morphology (6) and structure of the sperm, endanger the function of the sperm, and lead to male sterility (7). The quality standard for frozen bovine semen (GB4143-2008) stipulates that the proportion of morphologically abnormal spermatozoa in qualified bovine semen should be less than 18%, while the Real-World Health Organization Human Semen Analysis Laboratory Technical Manual (Fifth Edition) stipulates that the proportion of morphologically abnormal spermatozoa in human semen should be less than 4%. Therefore, understanding the regulatory mechanism of spermiogenesis and mining important functional genes have become the focus of exploration, prevention, and control of the causes of male breeding difficulty (8,9).

Therefore, we performed transcriptome analysis of round spermatids, elongated spermatids, and epididymal sperms of cattle. In order to identify differentially expressed genes that play an important role in spermiogenesis, the same stage mouse transcriptome sequencing data were introduced as a reference. Homology comparison analysis between species was used to obtain bovine genes with the same expression tendency in mice. Through PPI network, GO and KEGG pathway enrichment, and bioinformatic analysis of these genes, we initially obtained the protein interaction networks and pathways related to spermiogenesis, such as acrosomal and protamine replacement histones. At the same time, the structure and function of the important functional gene *ART3* were predicted, and the localisation of the ART3 protein in the seminiferous tubules was analysed by immunofluorescence. These results provide important clues and a theoretical basis for further exploration of the spermiogenesis mechanism, and play a positive role in improving the reproductive potential of cattle and promoting the exploration of human infertility mechanisms.

## Results

### Analysis of sequencing results of bull spermatids and sperm

After the quality control of the original sequencing data, the number of clean reads for each sample was more than 66.61 million, which accounted for more than 76.7% of the original sequencing data, and the Clean Q30 Bases ratio of each sample was higher than 91%, which showed that the sequencing data could be used in subsequent experiments (Table 1). After quality control, the data were compared with the bovine reference genome, the alignment ratio of clean reads was higher than 86% for each sample (Table 1), showing that the sequencing results could cover most of the reference genome, and further analyses could reveal the biological information of the genome during spermiogenesis.

**Table 1.** Sequencing results of bull spermatids and sperm

| Samples | R1 | R2 | R3 | E1 | E2 | E3 | M1 | M2 | M3 |
| --- | --- | --- | --- | --- | --- | --- | --- | --- | --- |
| Raw reads (million) | 76.983 | 94.627 | 78.303 | 78.018 | 77.196 | 78.670 | 77.515 | 76.957 | 88.457 |
| Clean Reads (million) | 72.947 | 72.556 | 71.185 | 73.697 | 71.376 | 71.570 | 66.614 | 71.147 | 79.564 |
| Clean Reads rate (%) | 94.8 | 76.7 | 90.9 | 94.5 | 92.5 | 91.0 | 85.9 | 92.5 | 90.0 |
| Clean Q30 Bases rate (%) | 94.0 | 93.1 | 93.2 | 94.3 | 94.0 | 93.8 | 91.0 | 92.6 | 92.6 |
| Mapped Reads (million) | 64.529 | 63.465 | 61.395 | 65.593 | 63.068 | 61.930 | 59.316 | 63.354 | 71.666 |
| Mapping Rate (%) | 88.4 | 87.5 | 86.2 | 89.0 | 88.4 | 86.5 | 89.0 | 89.0 | 90.0 |

Note: (1) Raw reads: The total number of sequences obtained from the original sequencing; (2) Clean reads: The number of reads of high quality sequence obtained after filtration; (3) Clean Reads rate: The ratio of the read number in the filtered high quality data to the read number in the original data; (4) Clean Q30 base rate (%): After filtration, the sequencing quality value of clean reads is more than 30, that is, the proportion of bases with an error rate is less than 0.1%. (5) Mapped reads : Number of clean reads sequences aligned to the reference genome. (6) Mapping Rate: Ratio of clean reads sequence number to reference genome. R: round spermatid; E: elongated spermatid; M: epididymal sperm. The same below.

### Principal component analysis (PCA) analysis of bull and mouse transcripts of spermatids and sperm

The PCA results in figure 2A show that the sum of PC1 (first principal component) (67.4%) and PC2 (second principal component) (19.8%) of bull reached 87.2%, of which round spermatid samples R1, R2, R3, elongated spermatid samples E1, E2, E3, and epididymal sperm samples M1, M2, and M3 were clustered together. The PCA results (figure 2B) showed that the sum of PC1 (83.9%) and PC2 (12.9%) of mice was 96.8%, among which round spermatid samples SRR3395024, SRR3395025, and SRR3395026, elongated spermatid samples SRR3395030, SRR3395031, SRR3395032, and epididymal sperm samples SRR4423201, SRR4423202, and SRR4423204 were clustered together.

**Figure 2.**
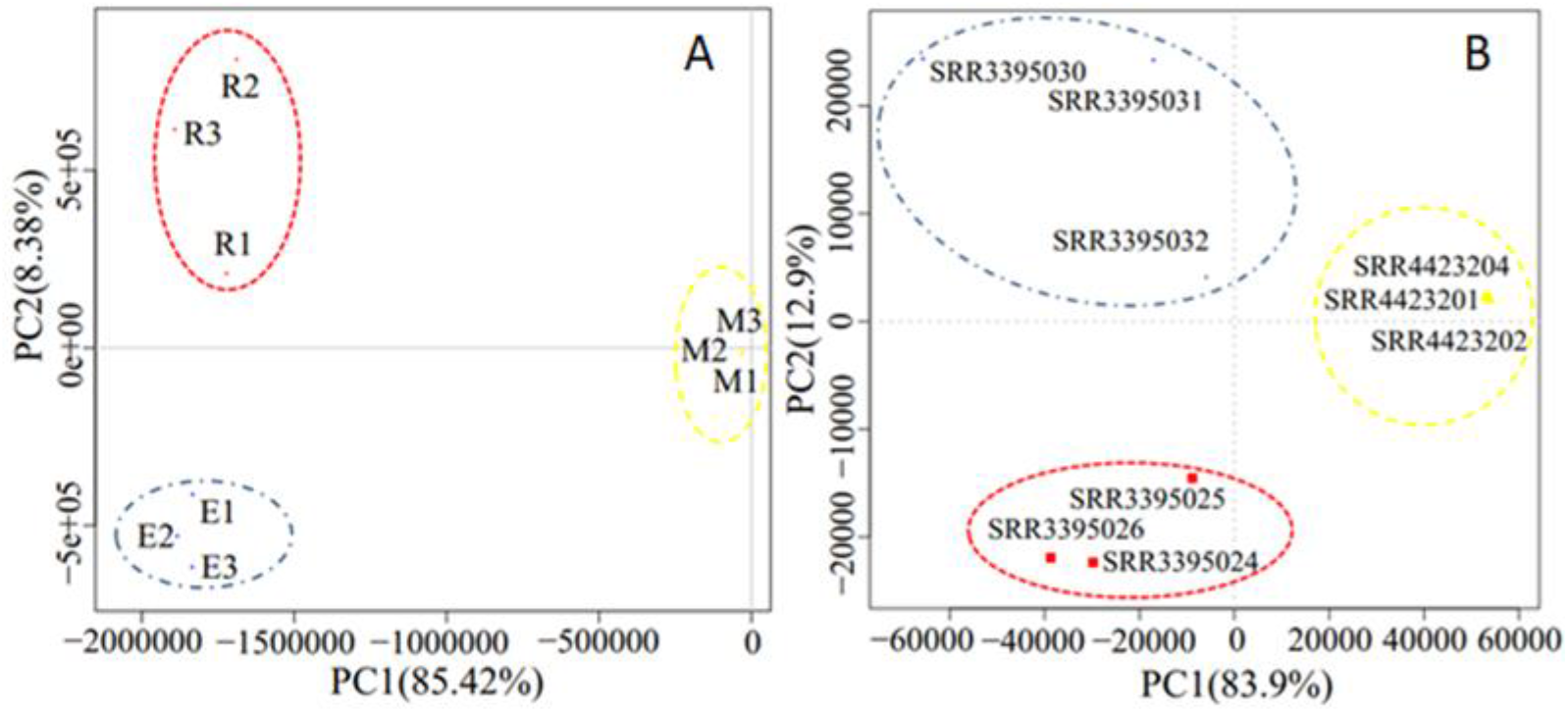
The bull (A) and mouse (B) PCA graphs of transcript expression of round spermatid, elongated spermatid and epididymal sperm. The red dotted circle represents round spermatids, the blue dotted circle represents elongated spermatids, and the yellow dotted circle represents epididymal sperms. The horizontal axis represents the first principal component (PC1), the horizontal axis represents the first principal component (PC2).

This showed that the transcripts of various types of spermatids from bulls and mice showed high intra-group repeatability, and the differences between the groups were large, so that the samples were representative.

### quantitative real-time PCR (qPCR) validation of the bull RNA-Seq results

The relative expression of 11 randomly selected genes was higher in round and elongated spermatids than in sperm, which was consistent with the variations from RNA-seq analysis (Fig.3). Among these genes, the relative expression levels of *LMTK2* and *MIGA2* showed no significant difference (*p* >0.05) in the sperm deformation stage; the relative expression level of *DNAL1* was significantly reduced from round spermatid to elongated spermatid (*p* < 0.05); *ART3, HIP1, SDHA*, and *YBX2* showed significant reduction in relative expression levels from elongated spermatid to epididymal spermatid (*p* <0.05). The relative expression levels of *TEKT2* and *PRKAR1A* were significantly reduced from both round spermatid to elongated spermatid and elongated spermatid to epididymal sperm (*p* <0.05); the relative expression level of *OAZ3* was significantly increased from round to elongated spermatid (*p* <0.05), and a significant decline from elongated spermatid to epididymal sperm (*p* <0.01). The qPCR results validated the RNA-Seq results, indicating that the bull transcriptome sequencing results were reliable, and subsequent experimental studies could be carried out.

**Figure 3.**
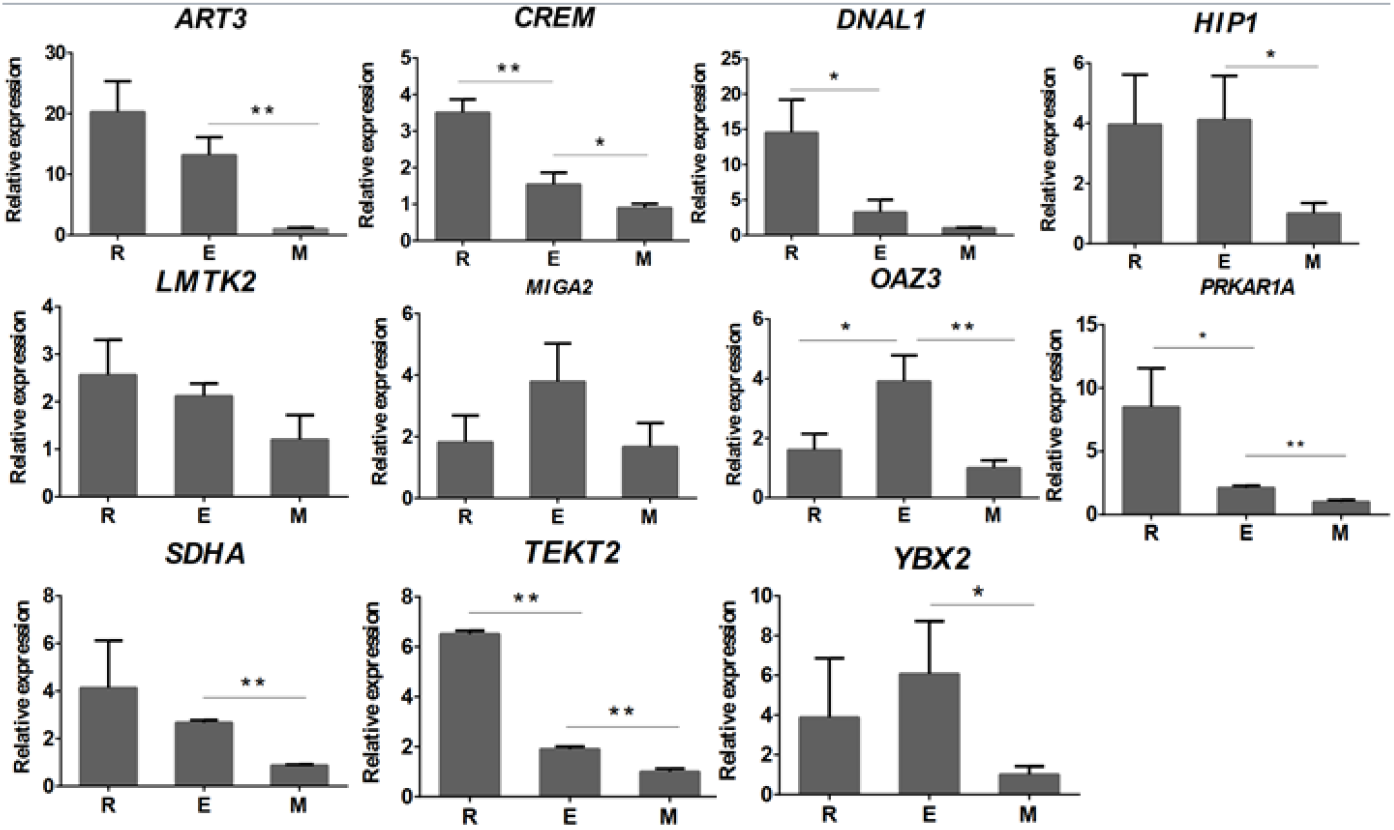
qPCR analysis of the expression levels of 11 genes in bull round spermatid, elongated spermatids, and epididymal sperm. Student’s t-test was used to assess the statistical significance of experimental data (\**p* < 0.05, \*\**p* < 0.01).

### Differentially expressed genes (DEGs) screening of the bull

A total of 7652 DEGs were obtained from the differential analysis of transcriptional expression of genes during spermiogenesis in the bull. Among these DEGs, 264 genes were upregulated and 253 genes were downregulated from round spermatids to elongated spermatids. The number of upregulated genes was slightly higher than that of downregulated genes. There were 241 upregulated and 7038 downregulated genes from elongated spermatids to epididymal sperm. The number of upregulated genes was far less than the number of downregulated genes. From the perspective of the entire spermiogenesis, the number of upregulated genes gradually decreased, and the downregulated genes gradually increased. In particular, at the later stage from the elongated spermatid to epididymal sperm, downregulated genes were greatly increased. A gradual decrease in gene expression was the main feature of this stage (Figure 4B, 4C).

**Figure 4.**
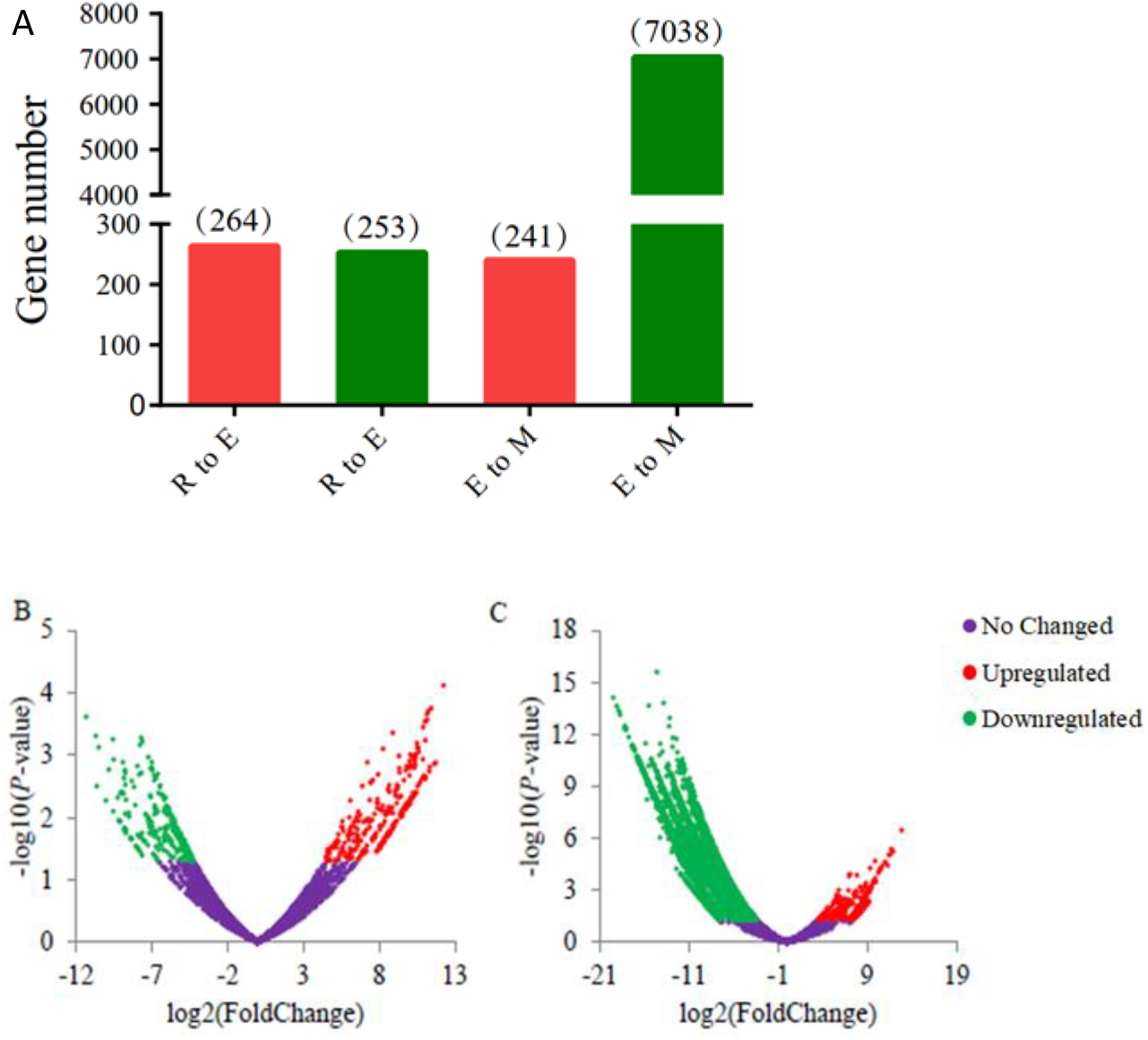
Statistical results of DEGs at each deformation stage of bull sperm. (A) Histogram of upregulated and downregulated genes during spermiogenesis. The abscissa R to E represents the stage of round to elongated spermatid, E to M represents the stage of elongated spermatid to epididymal sperm, and the ordinate represents the number of DEGs. (B) and (C) respectively represent DEGs volcano diagrams from round spermatid to elongated spermatid and from elongated spermatid to epididymal sperm. Among them, the abscissa represents the differential multiple of genes, and the ordinate represents the significance of the difference. *P*-value <0.05 and |log2 (Fold Change)|≥1 are used as the screening criteria for DEGs. Red indicates upregulated DEGs, green indicates downregulated DEGs, and purple indicates insignificant genes.

### Trend analysis of bull DEGs and mouse transcriptome homologous genes

Based on the 517 DEGs screened from round to elongated spermatids in bulls, a total of 344 homologous genes were found in the mouse sequencing data. Figure 5A and 5B illustrate that profile#0 had a statistically significant number of genes assigned (*p* <0.05), and they expressed a similar trend in both species. Gene expression decreased slowly in profile#0 during the whole sperm deformation stage, and there were 123 and 52 homologous genes in bull and mouse, respectively, and 33 overlapping genes (Figure 5C).

**Figure 5.**
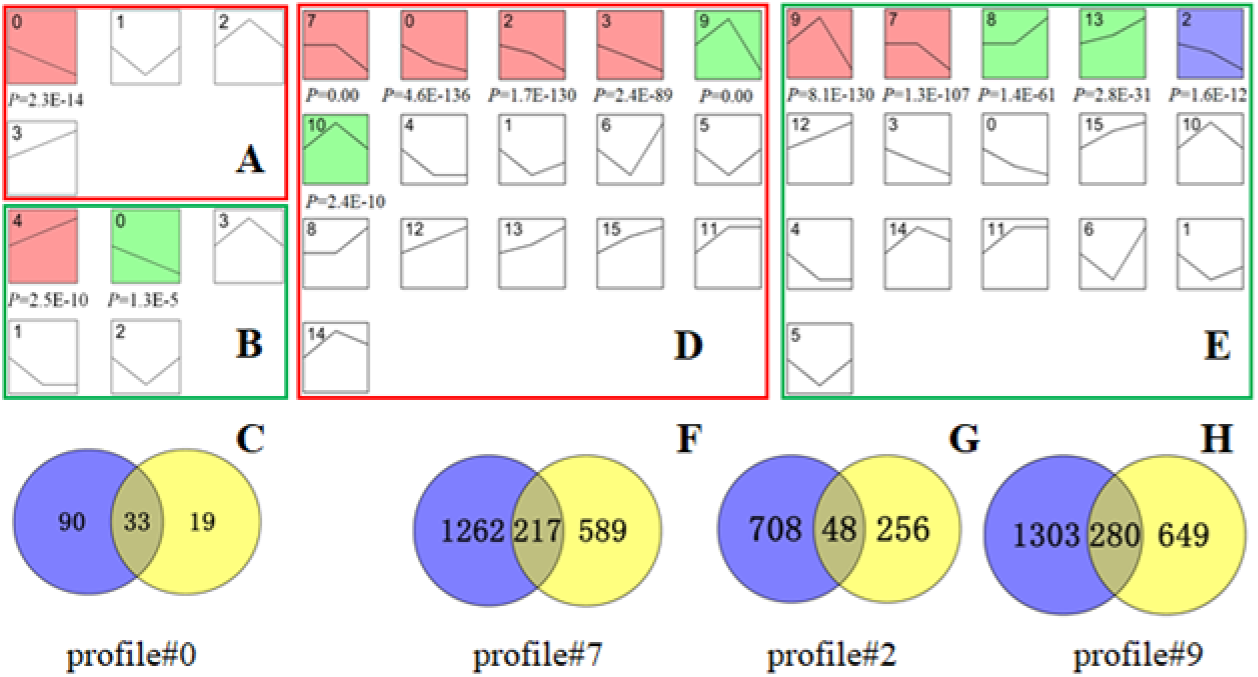
Screening results of DEGs with the same trend in bull and mouse. The bull (A, D) and mouse (B, E) trend chart of DEGs. Among them, diagrams A and B are trend diagrams of 344 homologous genes searched in mouse sequencing data for 517 DEGs at the stage of bull round to elongated spermatid. Diagrams D and E are trend diagrams of 6259 homologous genes searched in mouse sequencing data for 7279 DEGs from bull elongated spermatid to epididymal sperm. The colored profile graph indicates that the trend is statistically significant (*p* <0.05). After comparing figures A and B, the profile#0 gene set with the same expression trend and statistical significance was screened out. Image C is the Venn diagram of the cattle and mouse profile#0 gene sets. After comparing the D and E graphs, the profile#7, profile#2, and profile#9 gene sets with the same expression trend and statistical significance were screened. F, G, and H are Venn diagrams of the profile#7, profile#2, and profile#9 gene sets of bull and mouse, respectively. In figure C, F, G, and H, the blue gene set represents the bull homotopic gene and the yellow represents the mouse homotopic gene.

A total of 6259 homologous genes were found in the mouse sequencing data for the 7279 DEGs screened from the stage of elongated spermatid to epididymal sperm in bull. Figures 5D and 5E show that profiles #7, #2, and #9 had statistically significant numbers of genes assigned (*p* <0.05), and they were expressed in the same trends in the two species. Among them, the gene expression level from round to elongated spermatid was basically unchanged in profile#7, while it decreased slowly from elongated spermatid to epididymal sperm, the homologous genes for bovine and mouse were 1479 and 806, respectively, and 217 genes were overlapped; during spermiogenesis, the gene expression level in profile#2 gradually decreased, the homologous genes for bull and mouse were 756 and 304, respectively, and 48 genes were overlapped (Figure 5G). The expression level from round to elongated spermatid upregulated in profile#9, while the expression level from elongated spermatid to epididymal sperm was decreased, the homologous genes for bulls and mice were 1583 and 929, respectively, and 280 genes were overlapped (Figure 5H).

In summary, from the round to elongated spermatid stage, a total of 33 genes with the same homology and trend in bull and mouse were obtained. From the elongated spermatid to the epididymal sperm, a total of 545 genes with the same homology and trend in bull and mouse were obtained.

### GO and KEGG enrichment analysis of DEGs in bull homologous to mouse with same expression trend

The enrichment analysis results displayed that (Figure 6A), among the 33 genes, the genes in the terms of acrosomal vesicle (GO:0001669) and fertilisation (GO:0009566) might be involved in sperm acrosome formation; the genes in terms of sperm part (GO:0097223), reproduction (GO:0000003), and spermatogenesis (GO:0007283) might be involved in the deformation of the sperm head, tail, or other parts, thereby affecting sperm fertilisation ability; the genes in terms of ubiquitin-protein transferase activity (GO:0004842) might participate in the degradation of excess protein during sperm deformation through the protein-ubiquitinase system. The PPI analysis on DEGs selected in 3.5 from round to elongated spermatid in bull homologous to mouse with same trend showed that, *SPAM1, SPACA1, IZUMO1, TMEM190,* and *SPACA3* constituted the core regulatory network related to the formation of sperm acrosomes (Figure 6B).

**Figure 6.**
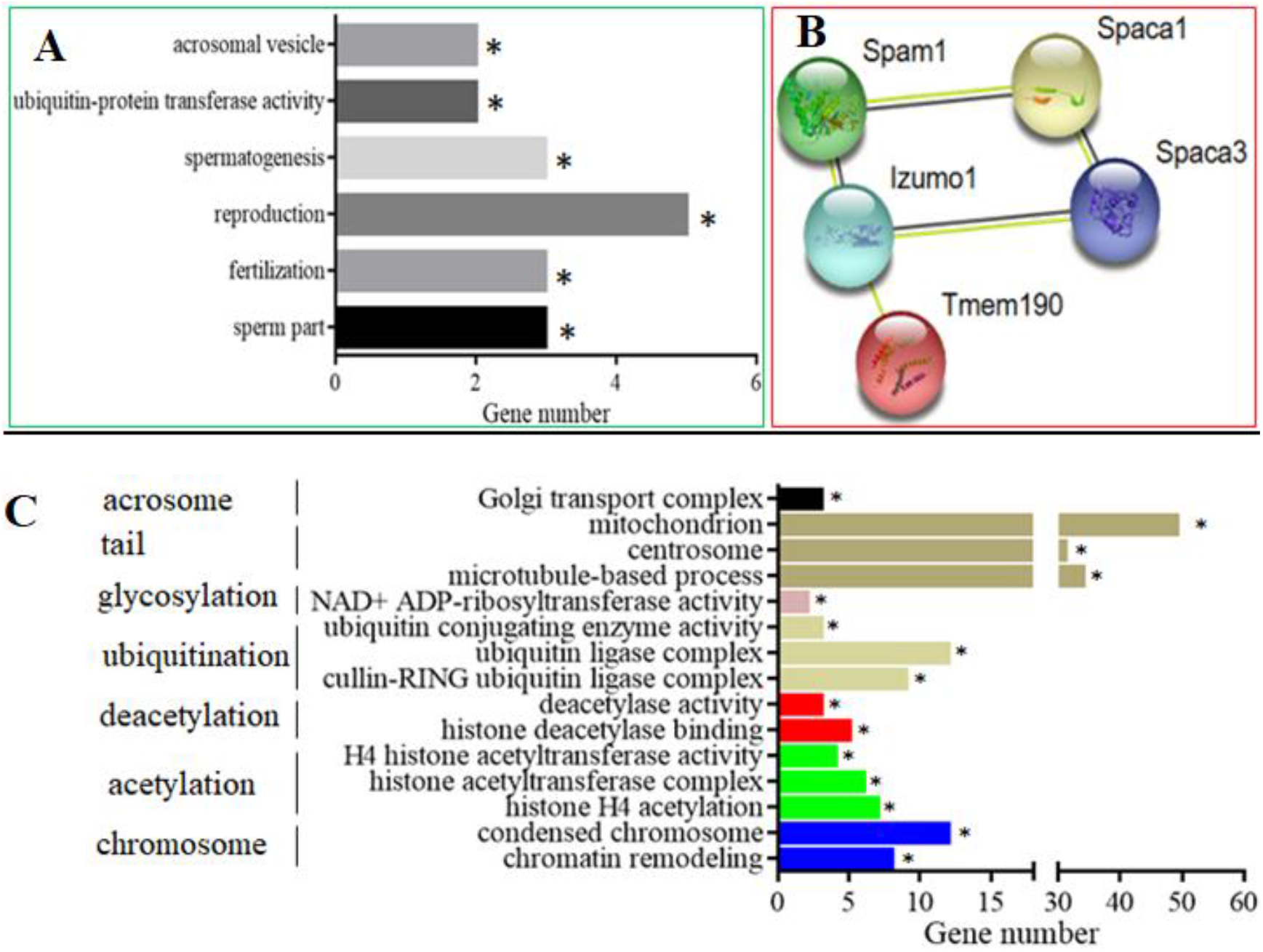
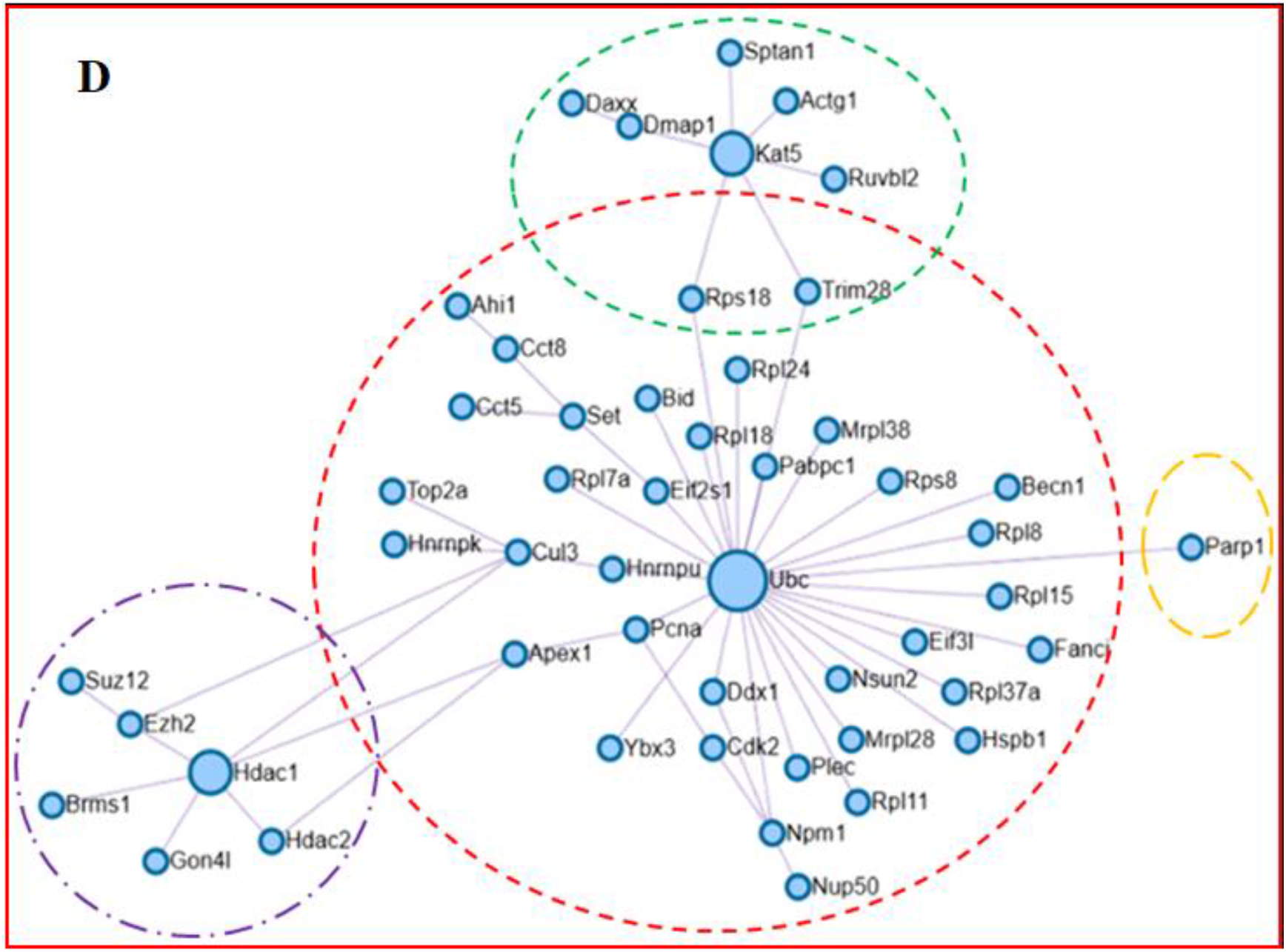
PPI and enrichment analysis results of the DEGs screened out from Figure 5. Figure A shows the enrichment terms of 33 DEGs screened in Figure 5C; figure B is the PPI regulatory network of the 33 DEGs, among which 5 genes constitute the core regulatory network related to sperm head formation. Figure C shows the enrichment terms of 545 DEGs screened in Figures 5F, 5G, and 5H; figure D is the PPI regulatory network of the 545 DEGs, among which 50 genes constitute the protein epigenetic modification regulatory core network. * indicates that the terms of gene enrichment are statistically significant (*p* ≤0.05).

The enrichment analysis results showed that (Figure 6C), among the 545 genes, the genes in the terms of Golgi transport complex (GO:0017119) might be involved in sperm head formation; the genes in the terms of mitochondrion (GO:0005739), centrosome (GO:0005813), and microtubule-based process (GO:0007017) might be involved in sperm tail formation. The genes in terms of ubiquitin-conjugating enzyme activity (GO:0061631), ubiquitin ligase complex (GO:0000151), and cullin-RING ubiquitin ligase complex (GO:0031461) might be involved in the degradation process of protein ubiquitination. The genes in terms of histone H4 acetylation (GO:0043967), histone acetyltransferase complex (GO:0000123), and H4 histone acetyltransferase activity (GO:0010485) might have acetylation activity, simultaneously, and the genes in the terms of histone deacetylase binding (GO:0042826) and deacetylase activity (GO:0019213) might have deacetylation activity. The balance of acetylation and deacetylation could play an important role in precisely regulating and controlling various pathways in cells through protein modification. The genes in the terms of condensed chromosome (GO:0000793) and chromatin remodelling (GO:0006338) might be involved in sperm chromosome remodelling, resulting in a gradual decrease in the transcription level. The PPI analysis of DEGs selected in 3.5 from elongated spermatids to epididymal sperm in bulls homologous to mice with the same trend showed that 50 genes constituted the core regulatory network related to the epigenetic modification of sperm histones (Figure 6D). In this network, the genes in the red dotted circle constituted a histone ubiquitination regulatory network with UBC as the core. The genes in the green dotted circle represent a histone acetylation regulatory network with *KAT5* as the core. The genes in the purple dotted circle represent a histone deacetylation regulatory network with *HDAC1* as the core. The *PARP1* gene in the orange dotted circle might repair DNA strand breaks caused by the replacement of histones by protamine through glycosylation.

Interestingly, the analysis found that PARP1 in the orange dotted circle in Figure 6D belonged to the NAD+ ADP-ribosyltransferase activity (GO:0003950), which is a large gene family. They might repair DNA strand breaks caused by the replacement of histones by protamine through glycosylation. The DEG *ART3* in this term might have the same effect as *PARP1*.

### ART3 immunohistochemical staining results

The immunofluorescence staining of ART3, a member of the ADP-ribosyltransferase family, is shown in figure 7. First, the staining results of protein ART3 in mouse sections are shown in figure 7A, 7B, and 7C. Both round and elongated spermatids in the seminiferous tubules showed red fluorescence signals of ART3 outside and the blue DAPI staining signals in the nuclei. However, only the blue DAPI staining signal could be detected in the other cells. Further analysis revealed that the distribution of fluorescence signals on the round and elongated spermatids of mice were polarised; in particular, the red signal was only distributed on the outer periphery of the spermatid in the lumen. Because the head of the elongated spermatid faces the basal lamina of the seminiferous tubule, and the tail faces the lumen, we speculated that protein ART3 might be related to the deformation and elongation of sperm cells, the formation of tails, or the shedding of cytoplasmic droplets. The staining results of protein ART3 in bull sections are shown in figure 7D, 7E, and 7F. The round spermatids and elongated spermatids still showed positive signals consistent with results from mice. However, the periphery of spermatocytes also showed a strong and uniform ART3 signal.

**Figure 7.**
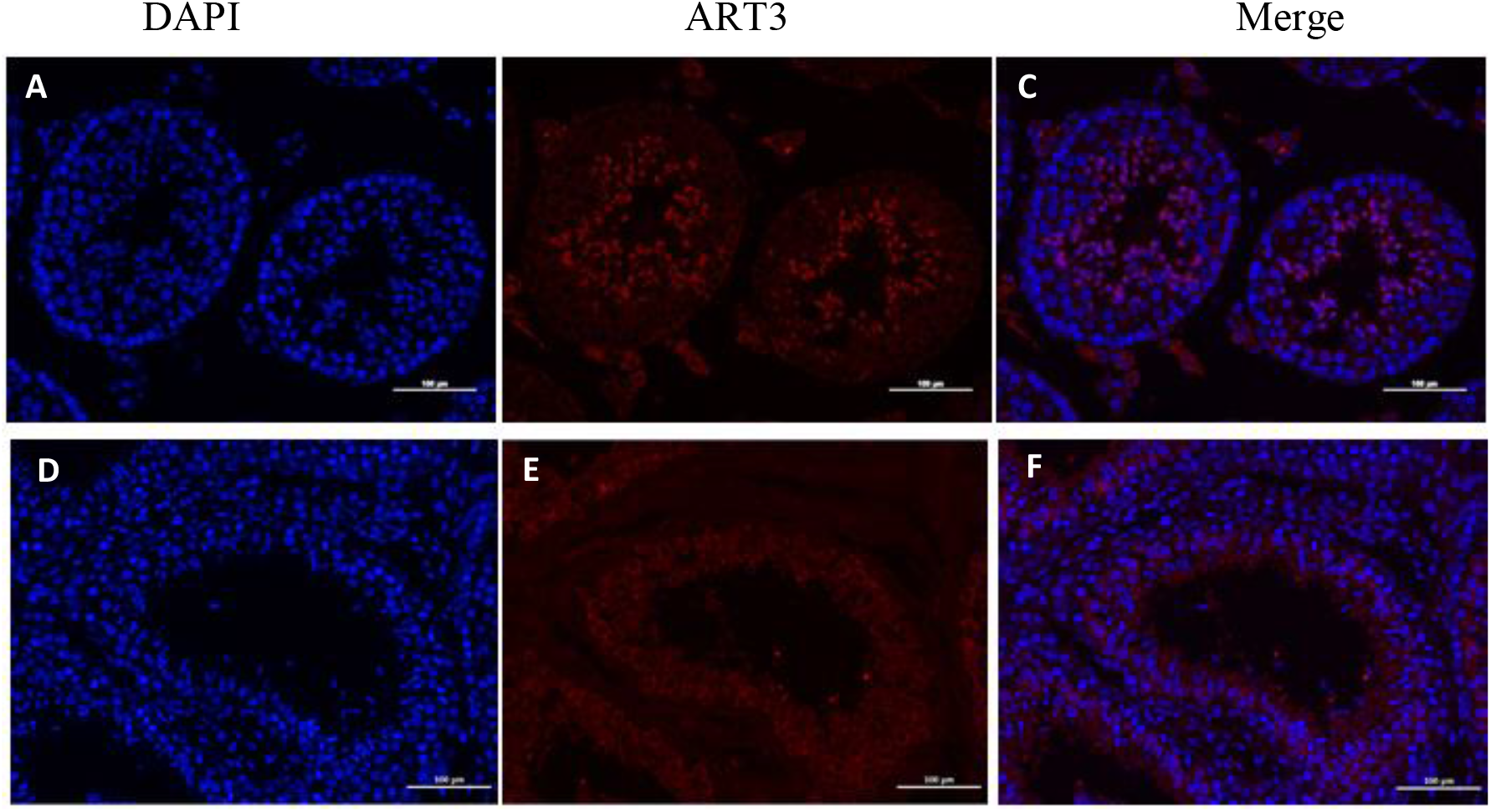
ART3 immunofluorescence staining. Figures A, B, and C show the staining results of cross sections of the seminiferous tubules of mouse testis, and Figures D, E, and F show the staining results of cross sections of the seminiferous tubules of bull testis. Blue is the nucleus colored by DAPI; red is the positive signal of ART3 antibody. C and F are merged images of ART3 and nuclear staining.

### Bioinformatic analysis of bovine ART3 protein

The bovine ART3 protein precursor (Q3T074) contained 390 amino acids, which were guided by the signal peptide into the Golgi apparatus and other subcellular structures after synthesis by the ribosome. The first 26 signal peptide sequence was degraded by signal peptidase, and a functional mature protein was secreted by the Golgi apparatus outside the cell membrane. The molecular weight of the mature protein ART3 was 41.071 kDa, the theoretical pI was 5.33, which was weak acid protein. Among its 20 amino acids, leucine (Leu) accounted for the highest proportion (8.79%), and tryptophan (Trp) accounted for the lowest proportion (0.55%); the molecular formula of the protein was C_1841_H_2839_N_473_O_554_S_19_, the half-life is 30 min (yeast, *in vivo*), and the instability index was 52.05, suggesting an unstable protein. The Hphob/Kyte & Doolittle value of 77% of the amino acids in the protein was less than 0, indicating that the protein was hydrophilic.

According to the prediction of transmembrane structure, the protein ART3 had no transmembrane region and was completely located outside the cell membrane with a signal peptide sequence in its precursor, it was speculated that it might be a secreted protein (PETITOT et al., 2016). The amino acid sequence of 56.04% was the random coil that played the role of connection in the secondary structure of the protein. Its N-terminal was mainly freely stretchable α-helices (26.65%), and the C-terminal was mainly a compact structure including extended strands and β-strands (17.3%). The structural characteristics of this polar protein with strong N-terminal flexibility and relatively compact C-terminal indicate that ART3 is likely to be an anchor protein (Mueller-Dieckmann et al., 2002).

Therefore, GPI anchor site prediction was performed, and omega-site position Pro was found at the C-terminus (position 359) of the ART3 protein. This site and the following hydrophobic amino acids constituted the conserved domain of the GPI-anchor protein, which further confirmed that protein ART3 was a GPI-anchor protein. The d1gxya template of the SCOPe2.06 database and single highest scoring were used to predict the tertiary structure of this anchored protein (Figure 8A). Based on its 215 consecutive amino acid residues with 100.0% credibility, the unique “four-stranded β-core” structure of the ART family was predicted, but the expected arginine-specific ADP-ribosyltransferases active site motif (R-S-EXE) was not found.

**Figure 8.**
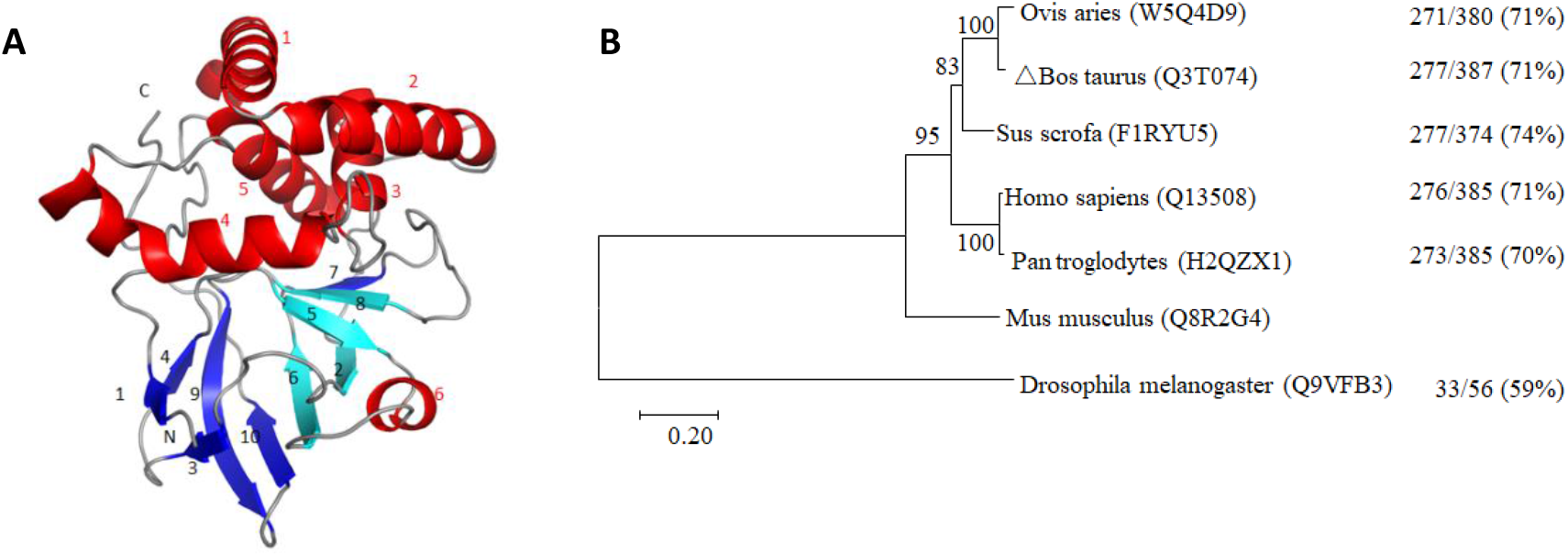
Tertiary structure (A) and phylogenetic tree (B) of bovine ART3 protein. The spirals, arrows, and loops in figure A represent the structures of α-helices, β-strands, and random coils, respectively. The light blue β2, β5, β6, and β8 indicate the “four-stranded β-core” conserved structure of the ART protein family. Figure B is a phylogenetic tree constructed by the Neighbor-joining method. The uniprot database number of each protein sequence is marked in brackets behind the species name. The branches in the horizontal direction represent the time changes in the evolution of the pedigree. 0.20 refers to the genetic variability of the ART3 protein. The value corresponding to the right side of each branch indicates the proportion of the full-length sequence of the same or similar site. The value corresponding to the right side of each branch indicates the proportion of the same sites plus similar sites of the ART3 protein of the species and the mouse to the full-length sequence.

In addition, the cluster analysis of the protein sequences among species in figure 8B showed that the *ART3* gene preceded the emergence of insects, such as *Drosophila*. The homology between them was 59%, which showed that the *ART3* gene plays an important role in the survival of organisms. During the long biological evolution, it showed higher species conservation and was not eliminated during natural selection. Its homology among higher mammals was even as high as 70%–74%. Moreover, clustering bovine and sheep into one category was consistent with their closer phylogenetic relationship, suggesting that the ART3 protein may play an important role in spermiogenesis.

## Discussion

Spermiogenesis is the result of the strict regulation of related genes expression in time and space (10). Analysis of the RNA-Seq data of bull and mouse showed an overall downward trend of gene expression during spermiogenesis. This is related to the successive replacement of histones by transition proteins and protamines during spermiogenesis and the gradual condensation of chromosomes. The haploid round spermatids are terminally differentiated cells and gradually become mature sperms, which only need to adapt to the environment to complete a major mission of fertilisation, and no longer have their own growth and development requirements (10,11). Therefore, scholars speculate that transcription does not occur during spermiogenesis (12,13), and internal transcripts are a residue from the spermatogenesis process (13,14). However, the results of this experiment showed that even though the transcription of more genes was downregulated, there were still upregulated genes from both round spermatids to elongated spermatids and elongated spermatids to epididymal sperm. Since approximately 15% of the DNA in the mature sperm is still bound to the histone, the transcription factor may have the opportunity to combine with the specific gene sequence to initiate gene transcription (15,16). Previous studies have shown that the transcriptional expression of *Gapds* in rat sperm was significantly increased in the late stage of round spermatids (17), and the transcriptional expression of genes such as presidents-cup was significantly upregulated from round spermatids into elongated spermatids in *Drosophila* (18). Therefore, with chromosome constriction during spermiogenesis, transcription gradually decreases, but it does not stop completely. An in-depth study of RNA-Seq data during spermiogenesis will help to reveal the physiological mechanism of spermiogenesis.

The mouse genetic background is explored deeply, and the introduction of homology and expression trend analyses of bovine and mouse RNA-Seq data will be conducive for revealing the physiological mechanism of bovine spermiogenesis, as well as the mining and identification of important functional genes (19,20).

Analysis of transcriptional DEGs from round to elongated spermatids showed that the DEGs mainly involved the formation of sperm acrosomes and fertilisation ability, which formed an interaction regulatory network with five key regulatory proteins. Among them, the membrane protein IZUMO1, located in the equatorial segment of the sperm head, may be the core regulatory gene in the network. Studies have shown that although Izumo^-/-^ mice can produce normal-shaped sperm that can pass through the zona pellucida, the sperm cannot fuse with the ovum (21). The membrane protein SPACA3 located in the acrosome can interact with the ovum plasma membrane oligosaccharide residue N-acetylglucosamine to allow the sperm to adhere to the ovum surface before fertilisation, in preparation for crossing the ovum plasma membrane (22). The acrosomal hyaluronidase SPAM1 enables the sperm to penetrate the hyaluronic acid-rich cumulus cell layer surrounding the oocyte (23). SPCA1 is located in the equatorial segment and other major parts of the sperm acrosome, play an important role in the formation of acrosome and sperm-oocyte fusion. SPCA1 deletion results in abnormal acrosome morphology and male mouse sterility (24). TMEM190, a regulatory protein of this network, not only localised on the inner acrosomal membrane of mouse, but also co-localises with IZUMO1 in the equatorial segment of the acrosome where the sperm-oocyte fused. However, co-immunoprecipitation experiments have shown that TMEM190 and IZUMO1 have no functional interaction, but TMEM190 deletion mice are fertile (25); therefore, its role TMEM190 plays in sperm-oocyte binding requires further experimental verification. In summary, this study found that the expression of genes involved in the regulatory network of acrosome formation showed a gradual downward trend during spermiogenesis, and reflected the acrosome formation mainly this stage.

The analysis of the DEGs from the elongated spermatid to epididymal sperm in this study showed that the replacement of histones by protamine is an important biological event at this stage. In order to ensure the occurrence of this biological event at an appropriate time, a regulatory network of histone acetylation centred on acetyltransferase KAT5 and a regulatory network of histone deacetylation centred on deacetyltransferase HDAC1 were formed to jointly serve the life activity of protamine replacement of histones. The expression level of *KAT5* in mouse testes is higher than that in other organs (26). It is located in the nuclear periphery near the round spermatid acrosome vesicle and elongated spermatids acrosome in the seminiferous tubules of the testis. Knockout of the KAT5 gene can destroy the hyperacetylation of histone H4, and the replacement of histones by protamine is blocked, resulting in abnormal sperm (27). Acetylation can reduce the affinity between nucleosomal histones and DNA, relaxing the chromatin structure (26). Histone hyperacetylation ensures that transition proteins and protamine replace histones in turn in elongated spermatids, the chromosomes are gradually densified, resulting in the downregulation of the transcriptional expression in sperm. The study revealed that the degree of histone hyperacetylation during spermatogenesis in infertile men was significantly lower than that in fertile men (28). The current transcriptomic analysis found that the expression of *KAT5* in round and elongated spermatids was higher than that in epididymal sperm, which was consistent with the replacement of histones with transition proteins and protamine during spermiogenesis, and also consistent with the downregulation of the transcriptional expression during spermiogenesis. Histone acetylation lays the structural foundation for the gradual replacement of histones. Histone acetylation also guarantees gene transcription in round spermatids (29). However, the degree of histone acetylation changes gradually in elongated spermatids, as the histones near the acrosome end are hyperacetylated, whereas the histones near the tail are only moderately acetylated (27). The biological significance of the change in the acetylation gradient requires further study.

The current study also found that the expression level of deacetyltransferase HDAC1 in round and elongated spermatids was higher than that in epididymal sperms. HDAC1 may act synergistically with KAT5 to ensure that protamine replaces histones at the appropriate time, which has been confirmed in mouse sperm studies (30). Studies have shown that HDAC1 is located in the nucleus in rat spermatocytes, round spermatids, and elongated spermatids, and can gradually stop transcription by promoting histone deacetylation (31). HDAC1 can prevent hyperacetylation of histones in the early round spermatid, to ensure that histones are replaced by protamines based on physiological requirements and normal and orderly spermiogenesis (32). The acetyltransferase activity increases in elongated spermatids, resulting in gradual hyperacetylation of histones, leading to its loose binding to DNA, which promotes the replacement of histones by protamines, and also leads to an increase in DNA fragmentation (33,34). Inhibition of HDAC1 expression can hinder spermatogenesis, reduce spermatogenesis, and lead to infertility (35).

The expression of HDAC1 at an appropriate time can regulate the timing and degree of hyperacetylation of histone H4 and avoid excessive DNA fragmentation (36). The deacetylation and acetylation regulatory networks cooperate, by regulating histone acetylation and deacetylation, to ensure the normal deformation of sperm.

We also found that the ubiquitin regulatory network centred on UBC (ubiquitin-conjugating enzymes, E2) played an important role from elongated spermatids to epididymal sperm. UBC binds ubiquitin molecules activated by ubiquitin-activating enzyme (UBA) E1 through thioester bonds, and then interacts with ubiquitin ligase enzymes (Ub ligase) E3 to transfer the ubiquitin molecules to the excess target protein residues or organelles to degrade it during spermiogenesis (37).

Histone ubiquitination can promote the replacement of histones by protamine and reparation of DNA strand breaks, which affects spermiogenesis (34), thereby affecting sperm morphology, vitality, and quantity (38). Therefore, some studies have proposed using ubiquitinase content to predict sperm fertilisation ability (39).

Ubiquitinated H2A and H2B appear in elongated spermatids, which can promote the removal and degradation of histones from chromatin (40). By combining with the regulation of acetylation and deacetylation, the histones in sperm can be replaced and degraded in an orderly manner, leading to a gradual decrease in transcription in sperm and the formation of a sperm head with dense, streamlined characteristics.

Our current study also found that *PARP1* plays an important role during spermiogenesis. This gene is located in the nucleus of spermatids and can repair DNA strand breaks and maintain sperm DNA integrity. *PARP1* deletion can prevent spermatids from maturation, apoptotic spermatids, and even male sterility (41). Under the action of the TOP2B enzyme, the DNA supercoil structure changes, and the electron leakage of mitochondria in spermatids produces excessive reactive oxygen species (ROS), which may cause DNA strand breaks (42). DNA strand breaks activate PARP1 within a few seconds, and cleave NAD+ into nicotinamide and ADP-ribose, and ADP-ribose polymerises into highly negatively charged poly(ADP-ribose) (PAR) to modify histones, repair broken DNA strands, and remodel chromatin structure (41).

Both ART3 and PARP1 belong to ADP-ribosyltransferases. This type of enzyme combines with the cholesterol-rich region of the cell membrane through a GPI-anchor (43), exposing the “four-stranded β-core” crack region composed of β2, β5, β6, and β8. After combining this region with NAD+ (44), the active site motif of this type of enzyme transfers the ADP-ribose on NAD+ to the amino acid residues of the target protein or polymerises ADP-ribose into PAR, while releasing the nicotinamide.

However, through bioinformatics analysis, we found that the bovine ART3 protein has no enzymatic active site motif, and consequently, cannot transfer ADP-ribose and cause ADP ribosylation of the target protein. In addition, HEK-293-T cell transfection (45) and DC27.10 cell overexpression (43) experiments showed that ART3 has no ADP-ribosyltransferase activity, which confirmed our bioinformatics analysis. ART3 has also been detected in human spermatocytes and is significantly related to human non-obstructive azoospermia (46). It is speculated that the transient expression of ART3 in human and bovine spermatocytes may facilitate the maturation of spermatocytes and provide a signal source for orderly cell division (45) to promote the development of spermatocytes to haploid spermatids. However, in the later spermatocytes in humans, ART3 may fall off due to the hydrolysis of the GPI-anchored structure (45,47).

While we detected ART3 protein in spermatocytes, round and elongated spermatids in bovines, its role in spermatogenesis had not been reported yet. Studies on cancer have shown that ART3 can be used as a triple-negative breast cancer (TNBC) marker, and overexpression in TNBC cells can reduce their apoptosis rate (48). Overexpression of ART3 in melanoma cells can promote cell migration (49). The continuous division of spermatogonia is similar to the immortal proliferation of cancer cells, and the migration of spermatogenic cells is similar to the metastasis of cancer cells. Based on these similarities, we speculated that genes expressed in spermiogenesis may also be similar to certain gene expression characteristics in cancer cells, namely tumor-testis gene expression patterns, such as PRAMEL1 (50), Xrcc1 (51), and TSLC1 (52). Immunofluorescence analysis in this study found that ART3 was located around round and elongated spermatids, and showed the polar positioning of ART3 toward the direction of sperm tail formation. Based on our study and the results of cancer research, it is speculated that ART3 is involved in sperm production, polarisation, and deformation, as well as the migration to the lumen of the seminiferous tubules and the shedding of droplets. However, further experimental verification is needed in the future.

In summary, we speculate that spermatocytes undergo meiosis and gradually differentiate into round spermatids under the induction of a signal source from ART3. Spermiogenesis begins from round spermatids, in which the formation of the acrosome is an important event, and the regulation network with IZUMO1 as the core gene is gradually formed. Among them, SPACA3 is necessary for the adhesion reaction between sperm and oocyte. SPAM1 promotes sperm to pass through the cumulus cell layer and finally cooperates with IZUMO1, SPACA1, and TMEM190 to ensure sperm-oocyte fusion. In the elongated spermatid stage, the important task of spermiogenesis is the densification of chromosomes and the formation of head and tail. The acetylase regulatory network with KAT5 as the core gene gradually hyperacetylated histones and loosens the structure between histones and DNA, creating conditions for transition proteins and protamines to replace histones.

At the same time, TOP2B changes the DNA supercoil structure, and histones H2A, H2B, and H3 are ubiquitinated, the replacement of protamine and the degradation of original histones are successfully realised. The broken DNA strands are repaired by PARP1 through glycosylation, causing the DNA strands to wrap around the smaller protamine. The deacetylation regulatory network with HDAC1 as the core gene ensures the occurrence of protamine substitution and histone degradation at an appropriate time, and finally finishes DNA constriction, leading to the formation of sperm heads with dense and streamlined characteristics. At the same time, ART3 promotes the polarization, deformation, and migration of spermatids. The formation of the sperm head and gradual decrease in gene expression are the most important events of spermiogenesis, which are achieved through the coordinated regulation of histone ubiquitination, acetylation, deacetylation, and ADP-ribosylation. In this study, the discovery of important regulatory networks and their related genes, such as acrosome formation, histone ubiquitination, acetylation, and deacetylation, as well as the discovery of important functional genes related to sperm polarity and tail formation, such as ADP-ribosylation, has unlocked the tip of the iceberg to further reveal the mechanism of spermiogenesis. It is of great significance to improve the reproductive potential of bovines and for diagnosing and treating male infertility.

## Experimental procedures

### Ethical statement

All animal breeding procedures and experiments were approved by the Animal Ethics Committee of the Institute of Animal Science, Chinese Academy of Agricultural Sciences (10 September 2019, CAAS; approval no, IAS2019-13).

### RNA-Seq analysis of the bull

Our previous research conducted a transcriptome analysis of the bull round and elongated spermatids and epididymal sperm; these data had three biological replicates for each cell type. Among them, round spermatids R1, R2, and R3 have 7698, 9463, and 78.30 million raw reads, respectively; elongated spermatids E1, E2, and E3 have 7802, 7720, and 78.67 million raw reads, respectively; and the epididymal sperm M1, M2, and M3 have 7751, 7696, and 88.46 million raw reads, respectively.

Raw reads from FASTQ were first processed using Perl scripts. Clean reads were obtained by removing the adaptor sequence from the reads, reads containing ploy-N, and low-quality reads from the raw data. Clean reads were aligned to the cattle reference genome (ARS-UCD1.2) using HISAT2 (v2.1.0) to build the index of the reference genome. StringTie (v2.1.2) assembled the alignments into full and partial transcripts, creating multiple isoforms as necessary and estimating the expression levels of all genes and transcripts (Pertea, 2016). DEGs of the three groups were analysed as previously described (Shi et al., 2017) using DESeq software (http://www.bioconductor.org/). The screening threshold to satisfy the conditions for a unigene would be defined as a DEG was a *p*-value < 0.05, and |log2 (fold-change)| ≥ 1.

### Principal component analysis (PCA) of transcripts among bull spermatids and sperms

We performed PCA using the dudi.pca function from the ade4 package in R. The data used for PCA were the transcript data of the bull round spermatid, elongated spermatids, and epididymal sperms.

### Validation of RNA-Seq data from the bull by qPCR

Three Holstein bulls with good health, 15 to 17 months old, similar body conditions, and normal fertility were selected to collect testes and epididymides. The collected testes and epididymides were placed in an ice box and quickly carried back to the laboratory. The epididymal sperms were collected using the float-out method (10). Diluting 1 μL of sperm sample with PBS and counted with a hemocytometer, and the sperm samples with an average viability of more than 50% in 3 visual fields were used for subsequent experiments. At the same time, the testis was cut into a tissue cube of about 1.0cm×0.6cm×0.4cm, and embed with optimal cutting temperature compound (OCT). It was placed in liquid nitrogen and frozen for 12-15 h, then transferred to a Leica CM1950 cryostat (Leica Microsystems; Wetzlar, Germany), which had been pre-cooled to −20 °C for 30 min, and then sliced to a thickness of 5 μm, filmed with a PEN2.0 membrane slide (RNase-free), and then stained with HistoGene LCM frozen section staining kit. The Leica LMD 7000 system was used to obtain round and elongated spermatids (Haiyang Zuo, 2016; Ren Xiaoxia, 2016; Li Xiaojun, 2018) and stored at −80 °C for future use.

Total RNA from round, elongated spermatids, and epididymal sperm samples was extracted with reference to the instruction manual of Trizol. cDNA was synthesised from 1 μg total RNA using a High Capacity cDNA Reverse Transcription Kit (Thermo Fisher Scientific, Waltham, MA, USA). Eleven genes were randomly selected 11 genes for qPCR verification, and all primers (Table 1) were designed using Primer 5. β-Actin was used to normalise the qPCR expression levels (10). The total reaction volume was 15μL, and contained 7.5 μL of 2×SG Green qPCR Mix (SinoGene, Beijing, China), 6 μL of nuclease-free water, 0.25 μL of forward primer (10 μM), 0.25 μL of reverse primer (10 μM), 1 μL of cDNA. qPCR was performed on a StepOnePLUS Sequence Detection System using the following parameters: initial denaturation at 95 °C for 10 min; denaturation at 95 °C for 20 s, annealing at 60 °C for 30s for 40 cycles, dissociation at 95 °C for 15 s, 60 °C for 30 s, and 95 °C for 15 s. 2^-ΔΔ^CT^^ method was used to analyse the expression level of genes.

### Acquisition and analysis of mouse RNA-seq data

In order to find functional genes with important value for life maintenance, homology analysis of bovine transcriptomic DEGs was conducted in mice with a relatively clear genetic background as a reference. Download RNA-seq data of mouse round spermatids, elongated spermatids, and epididymal sperms from the GEO database. Among them, the round spermatid sample numbers were SRR3395030, SRR3395031, and SRR3395032; elongated spermatid sample numbers are SRR3395024, SRR3395025, and SRR3395026; epididymal sperm sample numbers SRR4423201, SRR4423202, and SRR4423204. Construct an index file, create a full-length transcript, and calculate the transcript and gene expression.

### Trend analysis of gene expression and screening of genes with the same trend

Homologous comparison between bovine differentially expressed genes and mouse transcriptome was performed. Use STEM software (v1.3.12) (53) to perform trend analysis on the mouse transcripts and bovine DEGs, respectively. During the operation, the clustering method of the software was set to the STEM clustering method, and the other parameters were the default parameters. At the same time, the significance level was set at *p* <0.05. After the analysis of the results, we acquired a statistically significant profile that had the same expression trend in both bulls and mice. Then, the common genes of the two species were screened.

### GO term and KEGG pathway enrichment analysis

GO term and KEGG enrichment analysis of the selected common genes were performed using the webgestalt software (http://www.webgestalt.org/). During the operation, the enrichment analysis method of the software was set to the Over-Representation Analysis (ORA), and other parameters were the default parameters. GO terms and KEGG pathways with corrected P-values less than 0.05 were considered significantly enriched.

### Protein-protein interaction network (PPI network) analysis

Based on the BioGrid database (54), Metascape online software (http://metascape.org) was used to perform PPI analysis on the selected common genes to identify all the differentially expressed genes with related regulatory relationships. Then, we used the Molecular Complex Detection (MCODE) algorithm to screen modules with important biological functions in PPI (55).

### Immunohistochemical and Hematoxylin and Eosin (H&E) staining

The bull and mouse testis tissue cubes were soaked in 4% paraformaldehyde solution, fixed for 48 h, and then successively dehydrated in 70%, 85%, 95%, 100%, and 100% alcohol for 2 h. Then, the tissue cubes were successively immersed in a mixture of xylene and alcohol (1:1), xylene, and xylene in turn for 1 h. After transparent processing, the tissue cubes were embedded in paraffin and cut into 4 μm fine sections for subsequent immunohistochemical analysis and H&E staining.

Following deparaffinization and rehydration, the sections were subjected to heat-induced antigen retrieval by heating to 100°C in sodium citrate buffer (10 mM sodium citrate, 0.05% Tween 20, pH 6.0) for 10 min. After washing in PBS, the sections were blocked in 5% bovine serum albumin (BSA) for 45 min at room temperature (RT), and then incubated with anti-ART3 (1:100, sc-515157, Santa Cruz) overnight at 4°C. After incubation, slides were washed in PBS and then incubated with secondary anti-mouse antibody (1:500, B7389, Sigma) for 2 h at RT. Slides were washed with PBS and 3,3-diaminobenzidine (DAB) (Sigma, MA, USA) was added for 30 s, and then the slides were quickly washed in PBS. Slides were then counterstained in hematoxylin for 30 s before water washes and dehydration in ethanol and xylene. Slides were mounted with a coverslip. Images were captured using an Olympus BX 45 microscope with objective parameters of the UplanSApo 20×0.75 and 40x/0.95 objective. Representative 200x and 400x images are shown.

### Bioinformatics analysis of DEG ART3

The bovine ART3 protein precursor sequence was downloaded from the UniProt database (https://www.uniprot.org/), while the online server SignalP-5.0 Server was used to predict the protein signal peptide sequence and cut it off to obtain the mature ART3 protein sequence. The online server ExPASy-ProtParam (https://web.expasy.org/protparam/) was used to predict the physical and chemical properties of the mature ART3 protein sequence. The online server ExPasy-ProtScale (https://web.expasy.org/protscale/) was used to predict its hydrophilicity and hydrophobicity. At the same time, the ART3 protein structure was predicted. First, the online server TMHMM 2.0 Server (http://www.cbs.dtu.dk/services/TMHMM/) was used to predict its transmembrane structure, then using online servers SOPMA (https://npsa-prabi.ibcp.fr/cgi-bin/npsa_automat.pl?page=npsa_sopma.html) and Phyre2 (http://www.sbg.bio.ic.ac.uk/phyre2/html/page.cgi?id=index) predicted its secondary structure and tertiary structure, and the online server PredGPI (http://gpcr.biocomp.unibo.it/predgpi/pred.htm) was used to predict its GPI-anchor structure. Finally, MEGA7 (https://www.megasoftware.net/) software was used to construct a phylogenetic tree based on the ART3 protein sequence.

### Statistical analysis

All experiments were repeated at least three times. The differences between the treatment and control groups were analysed using one-way ANOVA or unpaired two-tailed t-test. *p*-values < 0.05 were considered to indicate statistically significant differences. All data were represented using the mean ±standard deviation (Mean ±SD). Analyses were performed using Microsoft Excel and GraphPad Prism 6.0 and R_x64 3.6.2.

## Data availability

Data will be shared upon request. Please contact Institute of Animal Science of CAAS

## Acknowledgments

We greatly thank Pro. Wang Dong and members of the lab for useful discussions. This research was funded by grants from the National Natural Science Foundation of China (31772595 and 31372296), for which we thank Wang Dong.

## Author contributions

D.W. conceptualization; D.W. data curation; X.L., C.Y.D. and R.Y.L. investigation; X.L. methodology; D.W. resources; D.W. supervision; D.W. funding acquisition; X.L., C.Y.D. and R.Y.L. validation; X.L. writing—original draft preparation; D.W. project administration; D.W., X.L., C.Y.D., R.Y.L. writing—review and editing

## Funding and additional information

This research was funded by grants from the National Natural Science Foundation of China (31772595 and 31372296). The content is solely the responsibility of the authors and does not necessarily represent the official views of the National Natural Science Foundation of China.

## Abbreviations

The abbreviations used are:
DEGs: Differentially expressed genes
GO: Gene ontology
KEGG: Kyoto Encyclopedia of Genes and Genomes
RT: Room temperature
qPCR: quantitative real-time PCR
ORA: Over-Representation Analysis
MCODE: Molecular Complex Detection
PPI: protein-protein interaction
ROS: reactive oxygen species
PCA: Principal component analysis

## References

1. Schenk, J. L. (2018) Review: Principles of maximizing bull semen production at genetic centers. Animal 12, 6

2. Fair, S., and Lonergan, P. (2018) Review: Understanding the causes of variation in reproductive wastage among bulls. Animal, 1–10

3. Schlegel, P. N. (2009) Evaluation of male infertility. Journal of the Louisiana State Medical Society Official Organ of the Louisiana State Medical Society 61, 261–283

4. Simon, L., Emery, B. R., and Carrell, D. T. (2017) Review: Diagnosis and impact of sperm DNA alterations in assisted reproduction. Best Practice & Research Clinical Obstetrics & Gynaecology 44, 38–56

5. Hamze, J. G., Sanchez, J. M., O’Callaghan, E., McDonald, M., Bermejo-Alvarez, P., Romar, R., Lonergan, P., and Jimenez-Movilla, M. (2020) JUNO protein coated beads: A potential tool to predict bovine sperm fertilizing ability. Theriogenology 155, 168–175

6. Devlin, D. J., Zaneveld, S. A., Nozawa, K., Han, X., Moye, A. R., Liang, Q., Harnish, J. M., Matzuk, M. M., and Chen, R. (2020) Knockout of mouse receptor accessory protein 6 leads to sperm function and morphology defects. Biol Reprod 102, 14

7. Evenson, D. P., and Wixon, R. (2008) Data analysis of two in vivo fertility studies using Sperm Chromatin Structure Assay-derived DNA fragmentation index vs. pregnancy outcome. Fertility & Sterility 90, 1229–1231

8. Neto, F. T. L., Bach, P. V., Najari, B. B., Li, P. S., and Goldstein, M. (2016) Spermatogenesis in humans and its affecting factors. Seminars in Cell & Developmental Biology, 10

9. Sironen, A., Lehti, and S., M. (2016) Formation and function of the manchette and flagellum during spermatogenesis. Reproduction: The official journal of the Society for the Study of Fertility

10. Zuo, H., Zhang, J., Zhang, L., Ren, X., and Wang, D. (2016) Transcriptomic Variation during Spermiogenesis in Mouse Germ Cells. Plos One 11, e0164874

11. Wang, Q., Lu, W., Yang, J., Jiang, L., Zhang, Q., Kan, X., and Yang, X. (2018) Comparative transcriptomics in three Passerida species provides insights into the evolution of avian mitochondrial complex I Comp Biochem Physiol Part D Genomics Proteomics 28, 10

12. D’Occhio, M. J., Hengstberger, K. J., and Johnston, S. D. (2007) Biology of sperm chromatin structure and relationship to male fertility and embryonic survival. Animal Reproduction Ence 101, 1–17

13. Ren, X., Chen, X., Wang, Z., and Wang, D. (2017) Is transcription in sperm stationary or dynamic? Journal of Reproduction and Development 63, 5

14. Fischer, B. E., Wasbrough, E., Meadows, L. A., Randlet, O., Dorus, S., Karr, T. L., and Russell, S. (2012) Conserved properties of Drosophila and human spermatozoal mRNA repertoires Proceedings of the Royal Society B: Biological Sciences 279, 2636–2644

15. Kramer, J. A., Mccarrey, J. R., Djakiew, D., and Krawetz, S. A. (2000) Human spermatogenesis as a model to examine gene potentiation. Molecular Reproduction & Development 56, 254–258

16. Wykes, S. M., and Krawetz, S. A. (2003) The Structural Organization of Sperm Chromatin. Journal of Biological Chemistry 2, 29471–29477

17. Welch, J. E., Barbee, R. R., Magyar, P. L., Bunch, D. O., and O’Brien, D. A. (2006) Expression of the spermatogenic cell-specific glyceraldehyde 3-phosphate dehydrogenase (GAPDS) in rat testis. Molecular Reproduction & Development 73, 1052–1060

18. Barreau, C., Benson, E., Gudmannsdottir, E., Newton, F., and White-Cooper, H. (2008) Post-meiotic transcription in Drosophila testes. Development 135, 7

19. Ghandhi, S. A., Sinha, A., Markatou, M., and Amundson, S. A. (2011) Time-series clustering of gene expression in irradiated and bystander fibroblasts: an application of FBPA clustering. Bmc Genomics 12, 1–23

20. Zheng, Y., Li, X., Huang, Y., Jia, L., and Li, W. (2018) Time series clustering of mRNA and lncRNA expression during osteogenic differentiation of periodontal ligament stem cells. Peerj 6, e5214

21. Inoue, Naokazu, Ikawa, Masahito, Isotani, Ayako, Okabe, and Masaru. (2005) The immunoglobulin superfamily protein Izumo is required for sperm to fuse with eggs. Nature

22. Ito, C., and Toshimori, K. (2016) Acrosome markers of human sperm. Anatomical Science International 91, 1–15

23. Andrée-Anne, Saindon, Pierre, and Leclerc. (2018) SPAM1 and PH-20 are two gene products expressed in bovine testis and present in sperm. Reproduction

24. Kishida, K., Harayama, H., Kimura, F., and Murakami, T. (2016) Individual differences in the distribution of sperm acrosome-associated 1 proteins among male patients of infertile couples; their possible impact on outcomes of conventional in vitro fertilization. Zygote 24, 654–661

25. Nishimura, H., Gupta, S., Myles, D. G., and Primakoff, P. (2011) Characterization of mouse sperm TMEM190, a small transmembrane protein with the trefoil domain: evidence for co-localization with IZUMO1 and complex formation with other sperm proteins. Reproduction 141, 437–451

26. Thomas, T., Loveland, K. L., and Voss, A. K. (2007) The genes coding for the MYST family histone acetyltransferases, Tip60 and Mof, are expressed at high levels during sperm development. Gene Expression Patterns 7, 657–665

27. Dong, Y., Isono, K. I., Ohbo, K., Endo, T. A., Ohara, O., Maekawa, M., Toyama, Y., Ito, C., Toshimori, K., and Helin, K. (2017) EPC1/TIP60-mediated histone acetylation facilitates spermiogenesis in mice. Molecular & Cellular Biology, MCB.00082-00017

28. Sonnack, V., Failing, K., Bergmann, M., and Steger, K. (2002) Expression of hyperacetylated histone H4 during normal and impaired human spermatogenesis. Andrologia 34, 7

29. Steger, K. (2001) Haploid spermatids exhibit translationally repressed mRNAs. Anatomy & Embryology 203, 323

30. Kim, J., Kim, J.-H., Jee, B.-C., Suh, C.-S., and Kim, S.-H. (2015) Is There a Link Between Expression Levels of Histone Deacetylase/Acetyltransferase in Mouse Sperm and Subsequent Blastocyst Development? Reproductive Sciences 22, 1387

31. Almabhouh, F. A., and Singh, H. J. (2018) Adverse effects of leptin on histone-to-protamine transition during spermatogenesis are prevented by melatonin in Sprague-Dawley rats. Andrologia 50

32. Hazzouri, M., Pivot-Pajot, C., Faure, A. K., Usson, Y., Pelletier, R., Sele, B., Khochbin, S., and Rousseaux, S. (2000) Regulated hyperacetylation of core histones during mouse spermatogenesis: involvement of histone deacetylases. Eur J Cell Biol 79, 950–960

33. Min, Y., Mi-Jeong, K., Sena, L., Eunyoung, C., and Ki-Young, L. (2018) Inhibition of TRAF6 ubiquitin-ligase activity by PRDX1 leads to inhibition of NFKB activation and autophagy activation. Autophagy 14, 15548627.15542018.11474995-

34. Kim, R. N., Kim, D.-W., Choi, S.-H., Chae, S.-H., Nam, S.-H., Kim, D.-W., Aeri Kim, A. K., Park, K.-H., Lee, Y. S., Hirai, M., Suzuki, Y., Sugano, S., Hashimoto, K., Kim, D.-S., and Park, H.-S. (2011) Major chimpanzee-specific structural changes in sperm development-associated genes. Funct Integr Genomics 11, 11

35. Betina, G., Gambini, P. C. R., H., S. M., D., V. A., Verónica, B., and R., G. C. (2018) Cocaine alters the mouse testicular epigenome with direct impact on histone acetylation and DNA methylation marks. Reproductive biomedicine online 37, 254–264

36. Gill-Sharma, M. K., Choudhuri, J., Ansari, M. A., and D’Souza4, S. (2012) Putative molecular mechanism underlying sperm chromatin remodelling is regulated by reproductive hormones. Clin Epigenetics 4, 20

37. Hanpude, P., Bhattacharya, S., Dey, A. K., and Maiti, T. K. (2015) Deubiquitinating enzymes in cellular signaling and disease regulation. Iubmb Life 67, 544–555

38. Sutovsky, and P. (2018) Review: Sperm–oocyte interactions and their implications for bull fertility, with emphasis on the ubiquitin–proteasome system. Animal, 1–12

39. Muratori, M., Marchiani, S., Criscuoli, L., Fuzzi, B., Tamburino, L., Dabizzi, S., Pucci, C., Evangelisti, P., Forti, G., and Noci, I. (2007) Biological meaning of ubiquitination and DNA fragmentation in human spermatozoa. Soc ReprodFertilSuppl 63, 153–158

40. Bao, J., and Bedford, M. T. (2016) Epigenetic regulation of the histone-to-protamine transition during Spermiogenesis. Reproduction 151, R55

41. Agarwal, A., Mahfouz, R. Z., Sharma, R. K., Sarkar, O., Mangrola, D., and Mathur, P. P. (2009) Potential biological role of poly (ADP-ribose) polymerase (PARP) in male gametes. Reprod Biol Endocrinol 7, 143

42. Meyer-Ficca, M. L., Lonchar, J. D., Ihara, M., Meistrich, M. L., Austin, C. A., and Meyer, R. G. (2011) Poly(ADP-Ribose) Polymerases PARP1 and PARP2 Modulate Topoisomerase II Beta (TOP2B) Function During Chromatin Condensation in Mouse Spermiogenesis. Biology of Reproduction 84, 900

43. Mario, L., Stephan, M., Rickmer, B., Bj?rn, R., Ann-Katrin, H., Kathrin, N., Lavinia, B., Peter, G., Hui, L., and Anna, Z. (2018) Proteomic Characterization of the Heart and Skeletal Muscle Reveals Widespread Arginine ADP-Ribosylation by the ARTC1 Ectoenzyme. Cell Reports 24, 1916–1929.e1915

44. Mueller-Dieckmann, C., Ritter, H., Haag, F., Koch-Nolte, F., and Schulz, G. E. (2002) Structure of the Ecto-ADP-ribosyl Transferase ART2.2 from Rat. Journal of Molecular Biology 322, 687–696

45. Friedrich, M., Grahnert, A., Paasch, U., Tannapfel, A., Koch-Nolte, F., and Hauschildt, S. (2006) Expression of toxin-related human mono-ADP-ribosyltransferase 3 in human testes. Asian Journal of Andrology 8, 281–287

46. Lin, F., Jiang, L., Yang, H., Yang, X., Wu, J., Huang, X., and Ni, W. (2015) Association of polymorphisms in ART3 gene with male infertility in the Chinese population. International Journal of Clinical & Experimental Medicine 8, 7944

47. Menzel, S., Rissiek, B., Bannas, P., Jakoby, T., Miksiewicz, M., Schwarz, N., Nissen, M., Haag, F., Tholey, A., and Koch-Nolte, F. (2015) Nucleotide-Induced Membrane-Proximal Proteolysis Controls the Substrate Specificity of T Cell Ecto-ADP-Ribosyltransferase ARTC2.2. Journal of Immunology 195, 2057–2066

48. Tan, L., Song, X., Sun, X., Wang, N., and Sun, Z. (2016) ART3 regulates triple-negative breast cancer cell function via activation of Akt and ERK pathways. Oncotarget 7

49. He, J., Li, Y., Wang, Y., Zhang, H., Ge, S., and Fan, X. (2018) Targeted silencing of the ADP-ribosyltransferase 3 gene inhibits the migration ability of melanoma cells. Oncology Letters

50. Mistry, B. V., Zhao, Y., Chang, T. C., Hiroshi, Y., Mitsuru, C., Jon, O., Francisco, D., Liu, W. S., and John, C. A. (2013) Differential Expression of PRAMEL1, a Cancer/Testis Antigen, during Spermatogenesis in the Mouse. Plos One 8, e60611-

51. Tebbs, R. S., Thompson, L. H., and Cleaver, J. E. (2003) Rescue of Xrcc1 knockout mouse embryo lethality by transgene-complementation. Dna Repair 2, 1405–1417

52. van der Weyden, L., Arends, M. J., Chausiaux, O. E., Ellis, P. J., Lange, U. C., Surani, M. A., Affara, N., Murakami, Y., Adams, D. J., and Bradley, A. (2006) Loss of TSLC1 causes male infertility due to a defect at the spermatid stage of spermatogenesis. Mol Cell Biol 26, 3595–3609

53. Ernst, J., and Bar-Joseph, Z. (2006) STEM: a tool for the analysis of short time series gene expression data. BMC Bioinformatics 7, 191

54. Stark, C., Breitkreutz, B. J., Reguly, T., Boucher, L., Breitkreutz, A., and Tyers, M. (2006) BioGRID: a general repository for interaction datasets. Nucleic Acids Res 34, D535–539

55. Bandettini, W. P., Kellman, P., Mancini, C., Booker, O. J., Vasu, S., Leung, S. W., Wilson, J. R., Shanbhag, S. M., Chen, M. Y., and Arai, A. E. (2012) MultiContrast Delayed Enhancement (MCODE) improves detection of subendocardial myocardial infarction by late gadolinium enhancement cardiovascular magnetic resonance: a clinical validation study. Journal of Cardiovascular Magnetic Resonance Official Journal of the Society for Cardiovascular Magnetic Resonance 14, 83–83

